# Adaptation of the *Romanomermis culicivorax* CCA-adding enzyme to miniaturized armless tRNA substrates

**DOI:** 10.1101/2020.07.07.190959

**Authors:** Oliver Hennig, Susanne Philipp, Sonja Bonin, Kévin Rollet, Tim Kolberg, Tina Jühling, Heike Betat, Claude Sauter, Mario Mörl

## Abstract

The mitochondrial genome of the nematode *Romanomermis culicivorax* encodes for miniaturized hairpin-like tRNA molecules that lack D- as well as T-arms, strongly deviating from the consensus cloverleaf. The single tRNA nucleotidyltransferase of this organism is fully active on armless tRNAs, while the human counterpart is not able to add a complete CCA-end. Transplanting single regions of the *Romanomermis* enzyme into the human counterpart, we identified a beta-turn element of the catalytic core that – when inserted into the human enzyme - confers full CCA-adding activity on armless tRNAs. This region, originally identified to position the 3’-end of the tRNA primer in the catalytic core, dramatically increases the enzyme’s substrate affinity. While conventional tRNA substrates bind to the enzyme by interactions with the T-arm, this is not possible in the case of armless tRNAs, and the strong contribution of the beta-turn compensates for an otherwise too weak interaction required for the addition of a complete CCA-terminus. This compensation demonstrates the remarkable evolutionary plasticity of the catalytic core elements of this enzyme to adapt to unconventional tRNA substrates.

## Introduction

tRNAs are the essential adaptor molecules which enable the decoding of the nucleic acid code into the amino acid sequence during the translational process (Rak *et al,* 2018). To fulfill this function, they need to undergo several maturation steps and interact with the translational machinery (Wolin & Matera, 1999; Hopper, 2013; Caetano-Anollés & Sun, 2014; Hopper & Nostramo, 2019). In higher organisms, this also includes the corresponding enzymes and proteins of mitochondria and chloroplasts (D’Souza & Minczuk, 2018; Zoschke & Bock, 2018). For the efficient recognition by a wide range of processing enzymes, translation factors as well as ribosomes, tRNAs fold into a conserved cloverleaf-like secondary structure consisting of acceptor stem, anticodon arm as well as D- and T-arm that adopts an equally conserved three-dimensional L-shape (Kim *et al,* 1974; Jühling *et al,* 2009; Giegé *et al,* 2012). The 3’- terminal CCA-triplet of the acceptor stem is a prerequisite for aminoacylation and the correct positioning of the charged tRNA in the ribosome (Sprinzl & Cramer, 1979; Green & Noller, 1997). In eukaryotes, this triplet is not genomically encoded but is added posttranscriptionally by tRNA nucleotidyltransferase (CCA-adding enzyme) (Weiner, 2004; Xiong & Steitz, 2006; Betat *et al,* 2010).

tRNA nucleotidyltransferases represent essential enzymes and are ubiquitously found in all domains of life. Representing members of the polymerase β superfamily, they split up into two classes, based on the composition of their catalytic core (Yue *et al,* 1996). Archaeal CCA-adding enzymes represent class I, while their bacterial and eukaryotic counterparts belong to class II (Yue *et al,* 1996). The overall sequence identity among both tRNA nucleotidyltransferase classes is rather low, although they catalyze the same reaction (Xiong *et al,* 2003). The catalytic core motif common in both classes consists of two aspartate residues DxD (x, any amino acids) that coordinate the catalytically important metal ions (Holm & Sander, 1995; Steitz, 1998; Aravind & Koonin, 1999). In class II enzymes, the DxD sequence belongs to motif A, one of the five conserved motifs A to E located in the N-terminal part of this tRNA nucleotidyltransferase type (Figure 1) (Li *et al,* 2002). Motif A catalyzes the nucleotide transfer onto the tRNA substrate via a two metal ion mechanism (Steitz, 1998). Motif B discriminates between ribose and deoxyribose (Li *et al,* 2002), while motif C is a flexible element which coordinates interdomain movements, contributing to the proper orientation of the substrates within the active center (Toh *et al,* 2009; Ernst *et al,* 2015). Motif D represents an amino acid-based template, where an arginine and an aspartate residue form Watson-Crick-like hydrogen bonds with the incoming nucleotide, and their orientation in the catalytic core determines the specificity for CTP or ATP, respectively (Li *et al,* 2002). Lastly, motif E is discussed to stabilize the positioning of the tRNA acceptor stem in the catalytic core (Li *et al,* 2002; Tomita *et al,* 2004).

**Figure 1.**
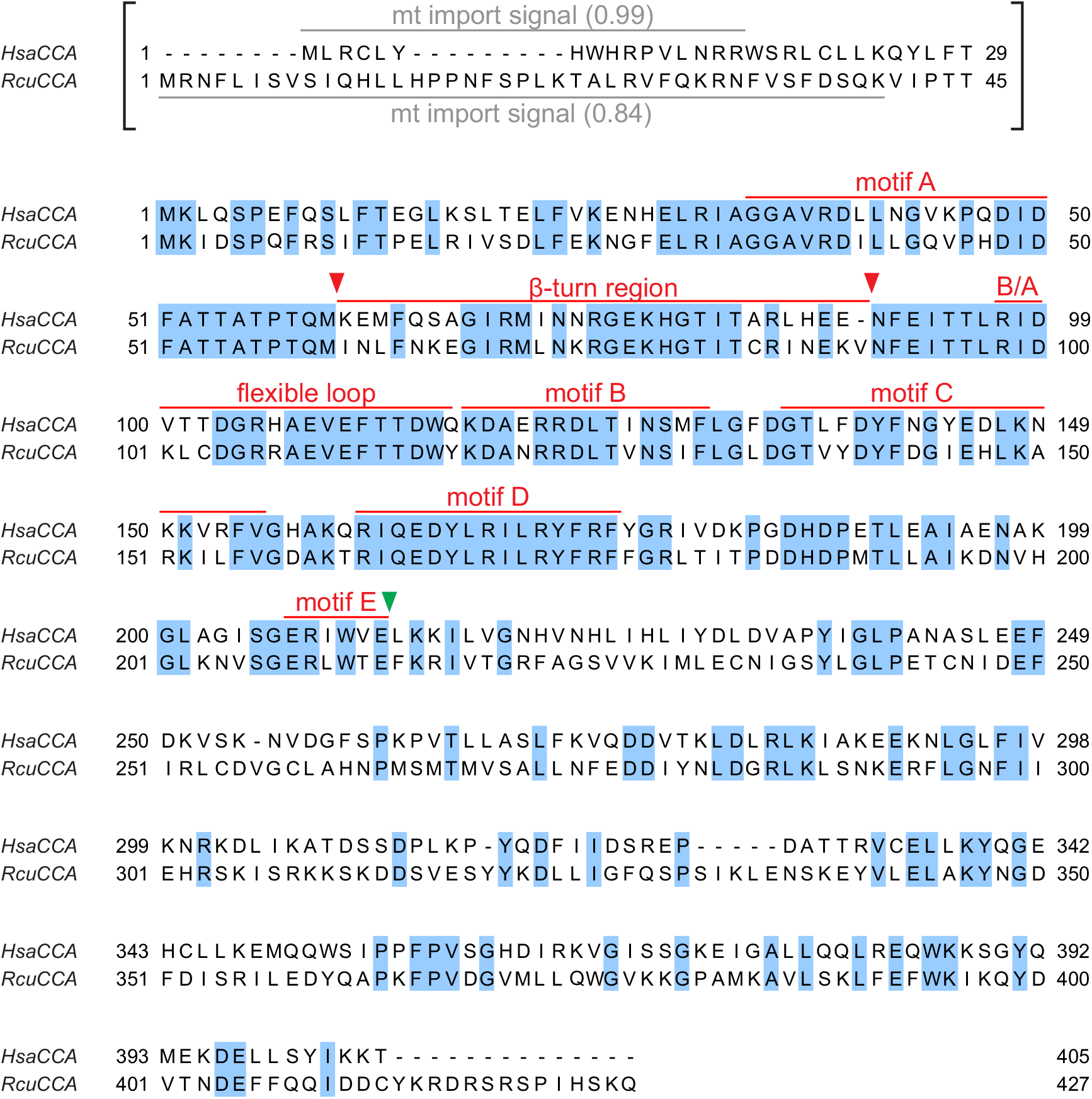
Sequence alignment of CCA-adding enzymes from *Romanomermis culicivorax* and *Homo sapiens*. Colored positions indicate identical residues. The N-terminal regions carrying the predicted mitochondrial import signals (grey bars, import probability is given in brackets) are shown in brackets and were excluded from the cloned open reading frames. Catalytically important elements (motifs A to E, β-turn element, basic/acidic motif B/A, and flexible loop) are labeled in red. Fusion position of reciprocal chimeras A and B are indicated by a green arrowhead (conserved position E212 (*Hsa*CCA) or E213 (*Rcu*CCA), located immediately downstream of motif E). Fusion positions of chimera H (β-turn element) are indicated by red arrowheads (K/I61 – E/V90).

Another catalytically important region is a flexible loop consisting of 10-20 amino acids that is located immediately upstream of motif B. While it is not conserved at the sequence level (Hoffmeier *et al,* 2010), its interaction with the amino acid template of motif D is required for the specificity switch from C to A incorporation, where it acts as a lever to accommodate the ATP in the nucleotide binding pocket (Neuenfeldt *et al,* 2008; Toh *et al,* 2009). Between motif A and the flexible loop, a β-turn element was identified that is also involved in A-addition, as it binds and positions the priming 3’-end of the growing CCA terminus in the catalytic core (Toh *et al,* 2009). Similar β-turn regions are present in many different polymerases, ranging from both classes of tRNA nucleotidyltransferases to poly(A) polymerases and DNA polymerases, underscoring the catalytically important function of this element (Davies *et al,* 1994; Sawaya *et al,* 1994; Cho *et al,* 2005; Tomita *et al,* 2006).

All these conserved motifs build up the active site in the N-terminal part of the enzyme. The C-terminus, in contrast, is much less conserved. Yet, it is of functional importance, as it is involved in tRNA binding, where it anchors the TΨC loop of the L-shaped tRNA substrate during nucleotide addition (Li *et al,* 1996; Tomita *et al,* 2004; Tretbar *et al,* 2011). Correspondingly, artificial CCA-adding substrates like mini- or microhelices are recognized and accepted for CCA incorporation at a much lower efficiency (Li *et al,* 1997; Lizano *et al,* 2008). Yet, metazoan mitochondria carry tRNA molecules that deviate from the cloverleaf structure, lacking either the D- or the T-arm (Wolstenholme *et al,* 1987; Okimoto & Wolstenholme, 1990; Watanabe *et al,* 2014). As an example, the mammalian mt-tRNA^Ser^(AGY) lacks the complete D-arm, so that D- and T-arm interactions do not exist (Bruijn *et al,* 1980; Hunter & Spremulli, 2004). In the mitochondrial genomes of nematodes, acari and arachnids, this situation comes to an extreme. These genomes are rich in genes for tRNAs that lack either the D- or the T-arm or even both (Wolstenholme *et al,* 1987; Watanabe *et al,* 1994; Klimov & Oconnor, 2009; Jühling *et al,* 2012; Pons *et al,* 2019). In the mermithid *Romanomermis culicivorax*, mt-tRNA molecules with the most dramatic truncations were identified, resulting in miniaturized hairpin-like tRNAs with a length of down to 45 nts, in contrast to the standard average tRNA size of 76 nts (Jühling *et al,* 2018; Wende *et al,* 2014). Such extremely truncated tRNAs fold into a three-dimensional boomerang-like shape that deviates from the consensus L-form (Jühling *et al,* 2018). Yet, these organisms encode for a single CCA-adding enzyme that has to act on both cytosolic as well as mitochondrial tRNA pools (Wolfe *et al,* 1994; Reichert *et al,* 2001; Nagaike *et al,* 2001; Tomari *et al,* 2002; Shikha & Schneider, 2020), and it was shown for the corresponding enzyme of *Caenorhabditis elegans* that it recognizes mt-tRNAs lacking D- or T-arm (Tomari *et al,* 2002). Since in *R. culicivorax* nine mt-tRNAs lack both arms, representing the strongest deviation from the consensus structure (Jühling *et al,* 2012), we investigated the co-evolution and substrate adaptation of its CCA-adding enzyme. In a comparative analysis, we identified the β-turn element as a major adaptation to the hairpin-like tRNA substrates. This adaptation is based on an increased substrate affinity of the enzyme. Hence, while the conventional substrate binding based on interactions between the enzyme’s C-terminus and the TΨC loop / T-arm is not possible with such tRNA hairpins, a different part of the enzyme took over this function to assure a sufficiently strong substrate binding for CCA-addition, demonstrating a surprising evolutionary plasticity of these enzymes.

## Results

### *Rcu*CCA adds a complete CCA-triplet to armless and canonical tRNAs

To investigate the catalytic activity and substrate specificity of the *R. culicivorax* CCA-adding enzyme (*Rcu*CCA), we identified a corresponding singular open reading frame in the *R. culicivorax* genome assembly available at https://parasite.wormbase.org/Romanomermis_culicivorax_prjeb1358/Info/Index/ (Schiffer *et al,* 2013). At the amino acid level, the overall sequence identity and similarity between this enzyme and the human tRNA nucleotidyltransferase (*Hsa*CCA) is 48 % and 66 %, respectively. Carrying the complete active site as well as a putative mitochondrial import sequence as predicted (Claros & Vincens, 1996), the N-terminus shows a correspondingly higher conservation (78 % sequence similarity; Figure 1). In the less conserved C-terminal part, *Rcu*CCA carries a short insertion of five residues and a terminal extension of 14 residues.

A conserved methionine residue downstream of the mt import signal was chosen as the N-terminus of the expressed open reading frame (labeled as position 1 in Figure 1). In previous experiments on the human enzyme, this position was successfully used and the absence of the mt import signal had no effect on its catalytic activity (Reichert *et al,* 2001; Augustin *et al,* 2003; Lizano *et al,* 2007; Lizano *et al,* 2008). The open reading frame was synthesized as a codon-optimized DNA sequence and recombinantly expressed in *E. coli*. Together with the corresponding enzymes from *E. coli* (*Eco*CCA; an organism exclusively carrying conventional cloverleaf-like tRNAs) and *H. sapiens* (*Hsa*CCA; an organism carrying conventional cytosolic as well as moderately reduced mt-tRNAs), the purified enzyme was tested *in vitro* for activity. As substrates, three different radioactively labeled tRNA transcripts were generated by *in vitro* transcription (Figure 2A), as it is well established that tRNA nucleotidyltransferases from all kingdoms readily accept *in vitro* transcripts lacking base modifications (Okabe *et al,* 2003; Hoffmeier *et al,* 2010; Ernst *et al,* 2018; Erber *et al,* 2020). tRNA^Phe^ from *Saccharomyces cerevisiae* is one of the best characterized tRNAs and represents a standard substrate for many tRNA-interacting enzymes (Oommen *et al,* 1992; Loria & Pan, 2000; Ernst *et al,* 2018), since the unmodified *in vitro* transcript folds into a structure almost identical to the native tRNA (Shi & Moore, 2000; Byrne *et al,* 2010).

**Figure 2.**
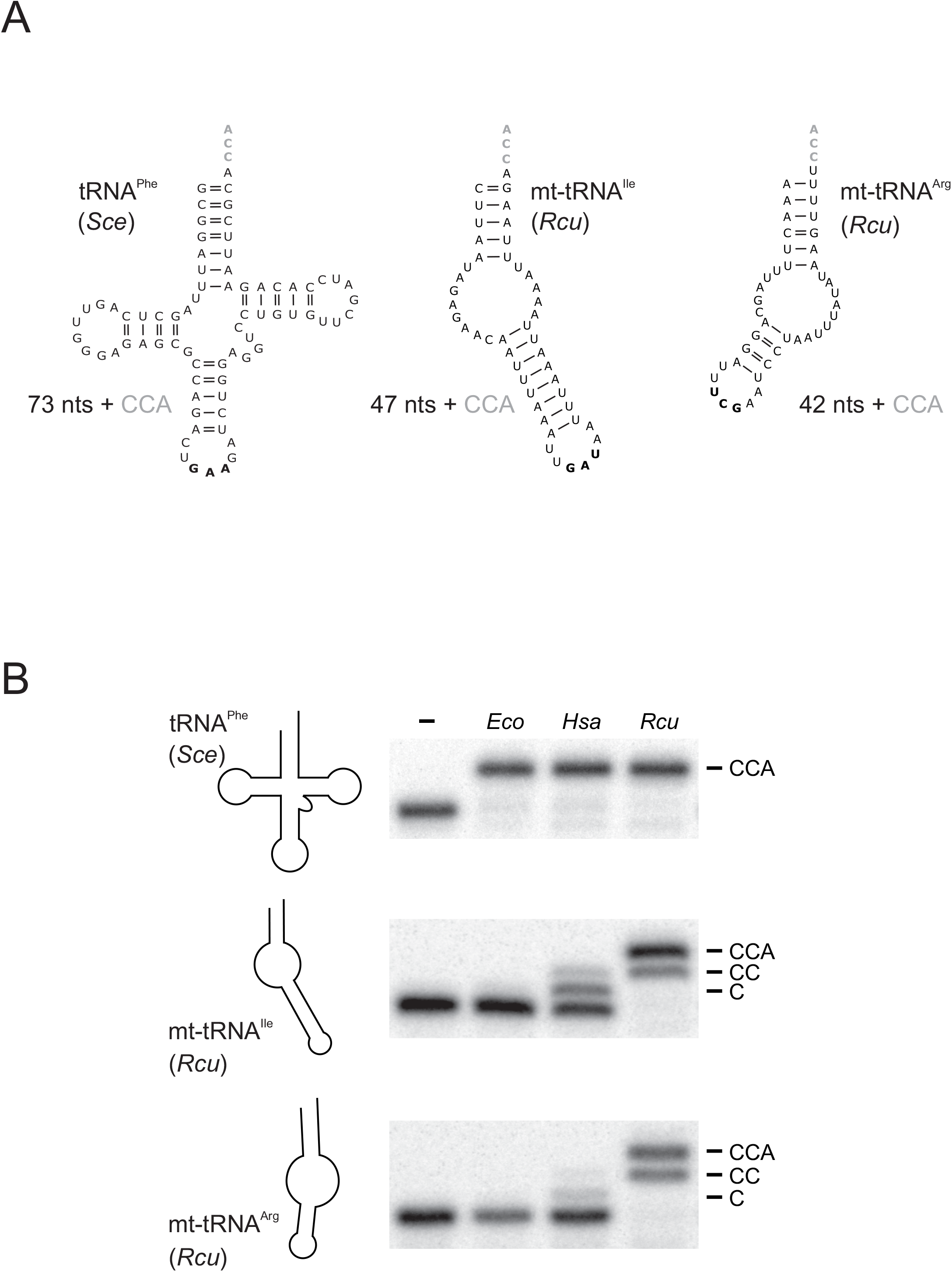
CCA-addition on conventional and hairpin-like tRNA substrates. **A.** tRNA^Phe^ from *S. cerevisiae* (*Sce*) represents a conventionally structured tRNA substrate of standard size (73 nts without CCA), while the mitochondrial tRNAs for isoleucine and arginine from *R. culicivorax* (*Rcu*) considerably deviate in size (47 and 42 nts, respectively; both without CCA) and structure, lacking both D- and T-arms. Anticodons are indicated in bold. **B.** CCA-addition on radioactively labeled tRNA transcripts catalyzed by the corresponding enzymes (20 ng each) of *E. coli* (*Eco*), *H. sapiens* (*Hsa*) and *R. culicivorax* (*Rcu*). Incubation without enzymes represent negative controls (-). All enzymes completely convert the canonical tRNA^Phe^ from *S. cerevisiae* into a mature transcript with CCA-end. On armless mt-tRNAs, the *E. coli* enzyme shows no activity at all, while the human enzyme adds only one or two C residues to mt-tRNA^Arg^ and mt-tRNA^Ile^, respectively. In contrast, the enzyme of *R. culicivorax* readily synthesizes a complete CCA-end on both transcripts, although the time of incubation was not sufficient for 100% A-addition. The experiment was done in three independent replicates. The panel shows a representative autoradiogram.

Furthermore, two armless mt-tRNAs from *R. culicivorax* were generated. With a length of 42 nts and 47 nts, respectively, mt-tRNA^Arg^ and mt-tRNA^Ile^ represent the shortest tRNAs identified so far, and the *in vitro* transcripts fold into hairpin-like structures with two single-stranded connector elements replacing D- and T-arm (Figure 2A) (Jühling *et al,* 2018). On the standard tRNA^Phe^, all enzymes added a complete CCA-triplet, indicating highly active enzyme preparations (Figure 2B). On the armless mt-tRNA substrates, however, the bacterial enzyme *Eco*CCA was completely inactive and did not add any nucleotides. The human enzyme *Hsa*CCA that has to recognize the human D-arm-lacking mt-tRNA^Ser^(AGY), catalyzed a moderate incorporation of two residues on the armless *Rcu* mt-tRNA^Ile^, but was almost inactive on *Rcu* mt-tRNA^Arg^. In contrast, *Rcu*CCA added complete CCA-triplets, regardless whether the substrate represented a conventional cloverleaf-structured tRNA or an armless hairpin-like tRNA, indicating an efficient adaptation to these miniaturized substrates (Figure 2B).

To investigate the substrate preferences of *Hsa*CCA and *Rcu*CCA in more detail, a Michaelis-Menten kinetics analysis was performed. Due to the limited RNA solubility, excessive saturating amounts of tRNA cannot be used, and the obtained parameters represent apparent values (Tomita *et al,* 2004; Tomita *et al,* 2006; Hoffmeier *et al,* 2010; Wende *et al,* 2015; Ernst *et al,* 2018). As *Eco*CCA showed no activity on armless tRNAs, this enzyme was excluded from further analysis. To discriminate between C- and A-addition, assays were performed on tRNAs lacking the CCA-end in the presence of either α-^32^P-CTP or α-^32^P-ATP and unlabeled CTP (Lizano *et al,* 2008; Just *et al,* 2008; Ernst *et al,* 2015). As shown in table 1, k_cat_ for CC-addition on tRNA^Phe^ is in a similar range for both enzymes, whereas the incorporation of the terminal A residue is somewhat less efficient for *Rcu*CCA (2-fold). On the armless tRNAs, the activity of *Hsa*CCA is strongly reduced when it comes to A-addition (Table 1, Figure 3A, C). While the enzyme adds the CC sequence at moderate efficiency on mt-tRNA^Ile^ and mt-tRNA^Arg^ (k_cat_ of 0.165 s^−1^ and 0.041 s^−1^ compared to 0.214 s^−1^ on tRNA^Phe^), its A-incorporation is strongly impaired on these substrates (k_cat_ of 0.006 s^−1^ on mt-tRNA^Ile^ and 0.003 s^−1^ on mt-tRNA^Arg^ versus 0.091 s^−1^ on tRNA^Phe^). In contrast, *Rcu*CCA readily accepts the armless tRNAs, although the overall efficiency is lower (Table 1, Figure 3B, D). Compared to the human enzyme, the corresponding k_cat_ values for CC-addition on all three types of tRNA are rather similar (tRNA^Phe^: 0.166 s^−1^ versus 0.214 s^−1^, mt-tRNA^Ile^: 0.224 s^−1^ versus 0.165 s^−1^, mt-tRNA^Arg^: 0.081 s^−1^ versus 0.041 s^−1^). However, A-addition on the armless substrates is much more efficient compared to *Hsa*CCA. *Rcu*CCA shows a k_cat_ of 0.052 s^−1^ (mt-tRNA^Ile^, comparable to the corresponding k_cat_ of 0.041 s^−1^ on tRNA^Phe^) and 0.012 s^−1^ (mt-tRNA^Arg^, 3.4-fold lower compared to tRNA^Phe^), while the human enzyme shows values of 0.006 s^−1^ (15-fold reduced compared to tRNA^Phe^) and 0.003 s^−1^ (30-fold reduced) on these substrates.

**Table 1.** Kinetic parameters of HsaCCA and RcuCCA for CC- and A-addition. Both enzymes show a rather similar efficiency in CCA-addition on a standard tRNA substrate. On armless tRNAs, however, the human enzyme is strongly affected in the addition of the terminal A residue. For each analysis, three independent experiments were performed.

| $k_{\text{cat}}$ [$\text{s}^{-1}$] | | *HsaCCA* | *RcuCCA* | relative activity<br>to *HsaCCA* |
| --- | --- | --- | --- | --- |
| tRNA<sup>Phe</sup><br>(*Sce*) | CCA* | 0.091 ± 0.012 | 0.041 ± 0.007 | ↓ 2 x |
|  | C*C* | 0.214 ± 0.034 | 0.166 ± 0.025 | ↓ 1.3 x |
| mt-tRNA<sup>Ile</sup><br>(*Rcu*) | CCA* | 0.006 ± 0.001 | **0.052 ± 0.011** | **↑ 8.7 x** |
|  | C*C* | 0.165 ± 0.016 | 0.224 ± 0.042 | ↑ 1.4 x |
| mt-tRNA<sup>Arg</sup><br>(*Rcu*) | CCA* | 0.003 ± 0.000 | 0.012 ± 0.001 | ↑ 4 x |
|  | C*C* | 0.041 ± 0.004 | 0.081 ± 0.011 | ↑ 2 x |

**Figure 3.**
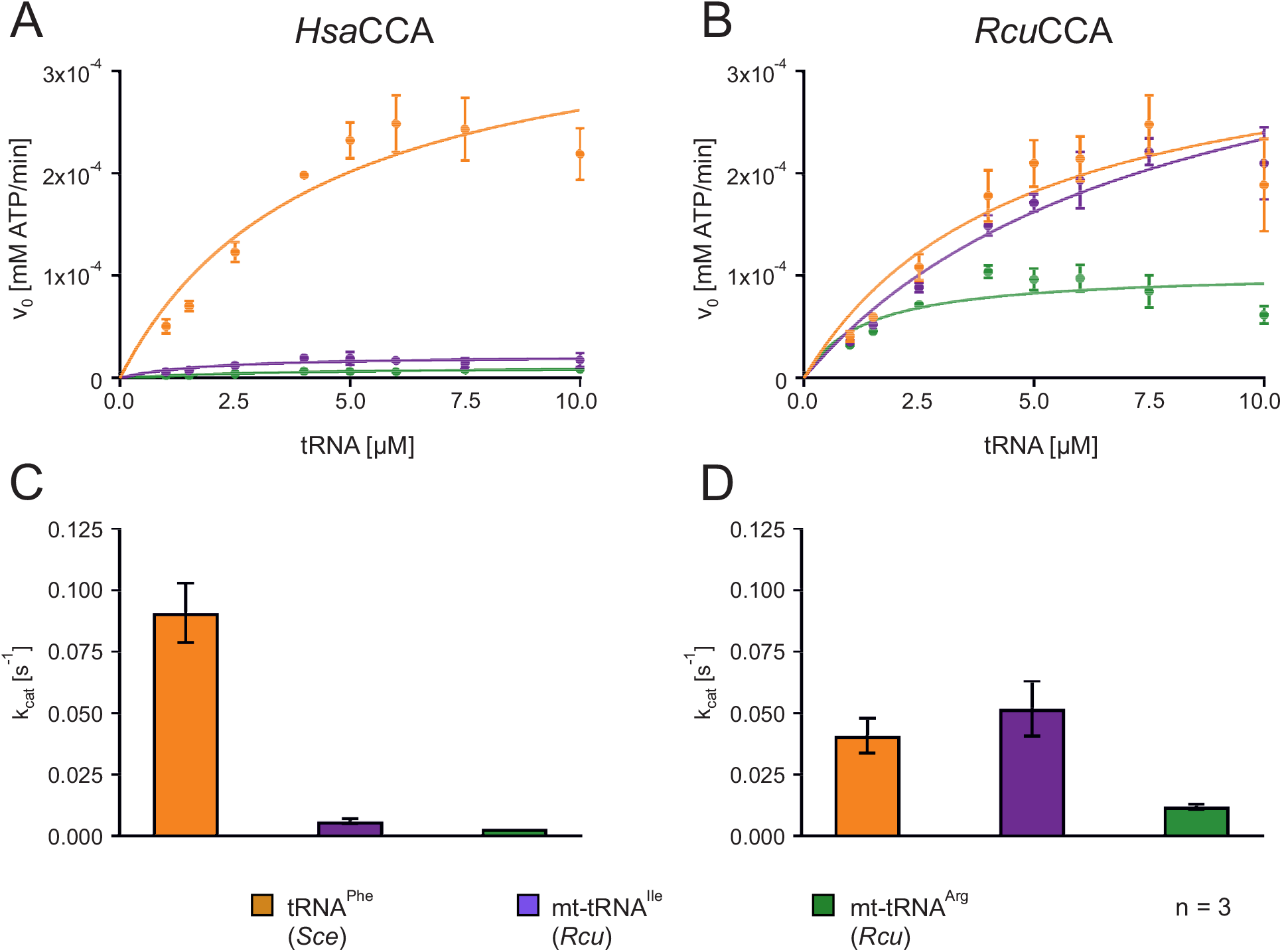
Kinetic analysis of CCA-addition. Reaction was monitored by the incorporation of radioactively labelled ATP. Curves representing turnover velocities are depicted for canonical tRNA^Phe^ (orange), mt tRNA^Ile^ (blue) and mt tRNA^Arg^ (green) with CCA-adding enzymes from *Homo sapiens* (*Hsa*CCA, **A**) or *Romanomermis culicivorax* (*Rcu*CCA, **B**). Error bars indicate standard deviation. Corresponding k_cat_ values were calculated by GraphPad Prism (**C**, **D**). *Hsa*CCA shows efficient processing of canonical tRNA^Phe^, but almost no activity on the armless mitochondrial tRNAs, while *Rcu*CCA readily accepts mt-tRNA^Ile^ as well as canonical tRNA^Phe^. On mt-tRNA^Arg^, *Rcu*CCA is also active in CCA-addition, but to a lesser extent.

Taken together, both enzymes prefer a conventionally structured tRNA as substrate. On the hairpin-like tRNAs, *Rcu*CCA still adds complete CCA-ends, although at a lower efficiency for mt-tRNA^Arg^ (Figure 3B). The human enzyme, however, strongly prefers cloverleaf-like tRNAs and is severely affected in A-addition on armless tRNAs, resulting in incomplete and hence non-functional tRNA molecules (Figure 3A).

### In the *Romanomermis* enzyme, especially the catalytic core is adapted to armless tRNA substrates

To identify the contribution of individual enzyme regions to the recognition of armless tRNAs as substrates for CCA-addition, we followed a strategy that we successfully applied to investigate several CCA-adding enzymes concerning their enzymatic features (Betat *et al,* 2004; Tretbar *et al,* 2011; Ernst *et al,* 2018). We reciprocally exchanged N- and C-termini of *Hsa*CCA and *Rcu*CCA, carrying the complete catalytic core and the region involved in tRNA binding, respectively. Based on the sequence alignment shown in Figure 1, we selected a glutamate at position 212 (*Hsa*CCA) and 213 (*Rcu*CCA), representing the last invariant residue of motif E (Martin & Keller, 2004) as fusion position. To allow for a direct comparison of enzymatic activities of the resulting proteins, we adjusted the efficiency of CCA-addition on the canonically structured tRNA^Phe^ substrate and defined an arbitrary unit as the amount of enzyme required for 50% substrate turnover, ranging between 0.3 ng for wild type (wt) enzymes and 0.5 to 1.3 ng for chimeras. Incubation of the substrate tRNA^Phe^ with 1 to 50 arbitrary units of both wt enzymes as well as chimera A (N-terminal catalytic core of *Rcu*CCA, C-terminus of *Hsa*CCA) and the reciprocal chimera B indicate that all enzymes are fully active and efficiently synthesize a complete CCA-terminus (Figure 4). On the minimalized mt-tRNA^Ile^, even 50 units of the human wt enzyme added only two C residues. In contrast, the same amount of *Romanomermis* enzyme added a complete CCA-end. Surprisingly, chimera B (with the catalytic core of *Rcu*CCA) synthesized a complete CCA-end at considerable efficiency, while identical units of chimera A, carrying the tRNA-binding C-terminus of *Rcu*CCA, catalyzed this reaction less efficiently.

**Figure 4.**
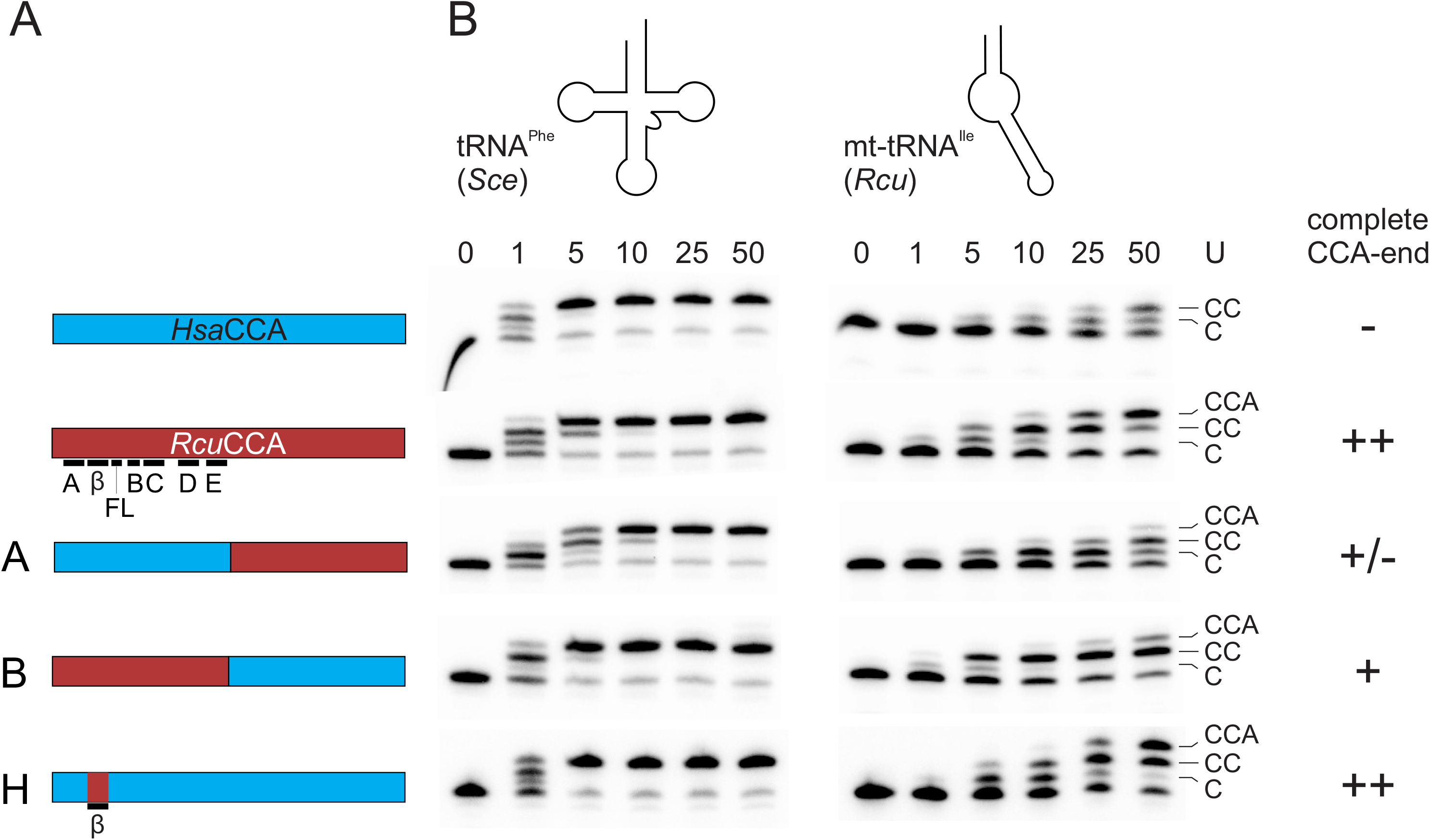
Catalytic activity of wild-type and chimeric enzymes on tRNA^Phe^ and mt-tRNA^Ile^. **A.** Bar representation of *Hsa*CCA (cyan) and *Rcu*CCA (red). Catalytic core elements are indicated in black. The reciprocal chimeras A and B are fused after position E213, downstream of motif E. The replaced β-turn element (β) in chimera H is located between motif A and the flexible loop (FL). **B.** CCA-addition on the conventional tRNA^Phe^ from yeast (*Sce*) and the armless mt-tRNA^Ile^ from *R. culicivorax* (*Rcu*) with increasing amounts of enzymes. All enzymes catalyze an efficient CCA-incorporation on the conventional tRNA. On the armless tRNA^Ile^, the *Romanomermis* wt enzyme synthesizes a complete CCA-end, while the corresponding human enzyme adds only two C-residues. Chimeras A and B also add the terminal A, but at a somewhat reduced level. On this tRNA substrate, chimera B, carrying the catalytic core of *Rcu*CCA, is more efficient than chimera A, where A-incorporation is only visible at the highest enzyme concentration. Chimera H shows an efficiency comparable to that of the *Rcu* wt enzyme, indicating the importance of the β-turn region in the reaction on armless tRNAs. The fact that chimera H is more active than chimera B likely reflects differences in the compatibility of the chosen fusion positions in these chimeras. For each construct, up to four independent experiments were performed. For each tRNA substrate, a representative gel is shown.

While these reactions indicate that both enzyme parts participate in the adaptation to the armless tRNA substrates, the contribution of the catalytic core seems to have a greater impact on the acceptance of these substrates. Based on these results, we generated a series of chimeras carrying parts of the *Rcu*CCA catalytic core in the context of the human enzyme (Figure EV1). With this “divide and conquer” strategy, we identified chimera H carrying the smallest catalytic core element of *Rcu*CCA, while it is still highly active on mt-tRNA^Ile^ (Figure 4). In this chimeric enzyme, a region of 30 amino acid residues (positions 61-91) is replaced that carries a β-turn element located between strands 3 and 4 of the β-sheet in the catalytic core (Martin & Keller, 2004), indicating that this region is involved in the adaptation of *Rcu*CCA to the armless tRNA substrates.

### The β-turn of the *R. culicivorax* CCA-adding enzyme strongly contributes to the interaction with armless tRNAs

To investigate how the individual regions of *Rcu*CCA participate in the recognition of armless tRNA substrates, we performed electrophoretic mobility shift experiments on an armless tRNA transcript and several enzyme chimeras which are able to catalyze the complete CCA-addition on hairpin-like tRNAs). To this end, radioactively labeled mt-tRNA^Ile^ lacking the CCA terminus was incubated with increasing amounts of recombinantly expressed enzymes and separated by native polyacrylamide gel electrophoresis (Figure 5). Enzyme-bound and free substrates were visualized and binding parameters were determined by nonlinear regression. As controls, wt enzymes *Hsa*CCA and *Rcu*CCA were included. The *Romanomermis* enzyme showed an efficient and robust binding to mt-tRNA^Ile^, resulting in a K_d_ value of 0.9 μM, while for the human enzyme, no significant binding could be detected, as previously reported for this enzyme class (Shi *et al,* 1998; Tretbar *et al,* 2011; Ernst *et al,* 2015). With K_d_ values of 1.5 and 1.4 μM, both *Hsa*/*Rcu* chimeras A and B show a binding behavior similar to that of the *Rcu*CCA wild type enzyme, demonstrating that both N- and C-termini contribute to the recognition of armless tRNAs. Surprisingly, chimera H, consisting of the human enzyme carrying the β-turn region of *Rcu*CCA, also showed a high affinity for mt-tRNA^Ile^ at a K_d_ of 1.8 μM.

**Figure 5.**
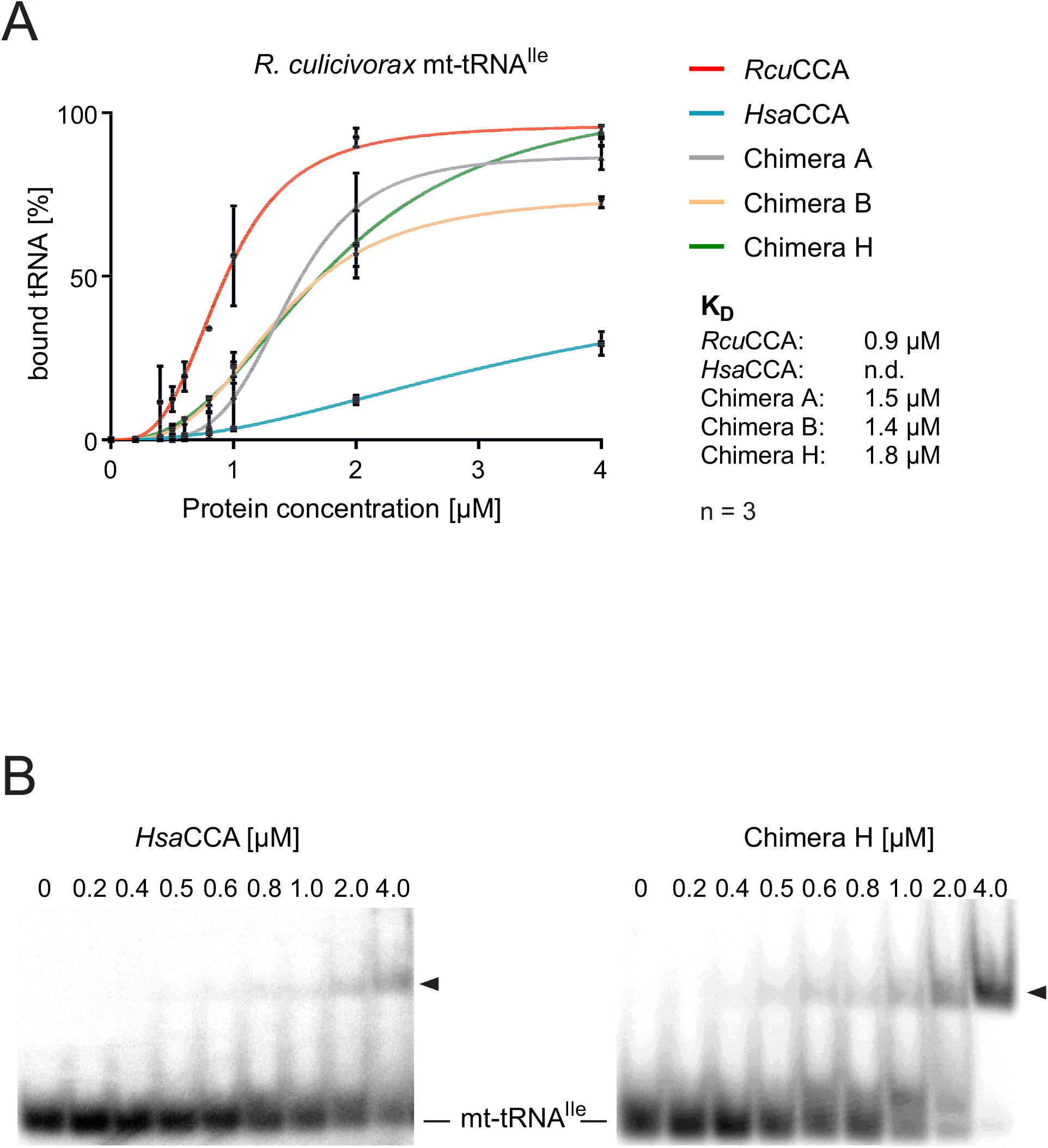
Binding of wt and chimeric CCA-adding enzymes to an armless tRNA. **A.** Quantitative analysis of enzyme binding to the armless mt-tRNA^Ile^ determined by electrophoretic mobility shifts. While the tRNA interaction of *Hsa*CCA over the whole concentration range (0 – 4 μM) is too weak to calculate dissociation constants, *Rcu*CCA as well as chimeras A, B and H exhibit a strong affinity to this substrate, resulting in dissociation constants in the range of 0.9 to 1.8 μM, respectively. Data are means ± SD; n = 3. **B.** Representative example of shifted tRNA – enzyme complexes. The human enzyme shows only very weak and inefficient binding to the armless mt-tRNA^Ile^ (left panel). In contrast, chimera H, where the β-turn element in the human enzyme is replaced by the corresponding region of *Rcu*CCA, exhibits an efficient substrate interaction that results in a dissociation constant of 1.5 μM (right panel). Shifted bands are indicated by arrow heads.

Gel shift experiments on the conventional substrate tRNA^Phe^ indicate that this high affinity of the *Rcu*CCA enzyme is not restricted to armless tRNA substrates (Figure EV2). Rather, the enzyme exhibits a similar binding behavior to the cloverleaf-shaped tRNA, while the human enzyme again shows almost no interaction. Hence, the adaptation of the *Romanomermis* CCA-adding enzyme to process both cloverleaf-structured cytosolic as well as armless mitochondrial tRNAs is obviously achieved by a general exceptional tight interaction with its substrates, regardless whether they represent canonical or minimalized tRNAs.

## Discussion

### A specific adaptation in the catalytic core enables CCA-addition to minimalized tRNA substrates

As the tRNA genes of most organisms do not encode the 3’-terminal CCA-triplet, this essential feature has to be added posttranscriptionally by the CCA-adding enzyme. In eukaryotes, a single enzyme is responsible for this maturation step in both cytosolic as well as mitochondrial tRNA pools (Chen *et al,* 1992; Reichert *et al,* 2001; Nagaike *et al,* 2001). Similar to other general tRNA maturation enzymes, the CCA-adding enzyme recognizes not a specific sequence or base pair in its substrates, but relies on common elements of the overall tRNA cloverleaf structure, as it is also observed for RNase P, tRNase Z and some tRNA modifying enzymes (Levinger *et al,* 1998; Zahler *et al,* 2003; Kirsebom, 2007; Levinger *et al,* 2009; McKenney *et al,* 2017). Metazoan mitochondria, however, encode for tRNAs with deviations from the cloverleaf, where D- or T-arms are reduced or lacking (Bruijn *et al,* 1980; Wolstenholme *et al,* 1987; Wende *et al,* 2014). As these tRNAs still carry a conventional acceptor stem, they are correctly processed by RNase P, since this enzyme predominantly recognizes this structural feature (Zahler *et al,* 2003; Kirsebom, 2007). The CCA-adding enzyme, in contrast, specifically interacts with the tRNA elbow region, and especially with the T-loop region (Spacciapoli *et al,* 1989; Spacciapoli & Thurlow, 1990; Hegg & Thurlow, 1990; Li *et al,* 1996; Shi *et al,* 1998). Hence, a bacterial enzyme like the *E. coli* version, evolved for conventional tRNAs, is strongly impaired on truncated tRNAs (Tomari *et al,* 2002). The most extreme truncations are observed in nematodes like *R. culicivorax*, in acari and in arachnids, where hairpin-like tRNAs are found that require a specific co-evolution of the corresponding enzymes. On such substrates, the *E. coli* enzyme is completely inactive (Figure 2B). The human enzyme, however, co-evolved to accept the D-armless tRNA^Ser^_AGY_ found in human mitochondria (Bruijn *et al,* 1980). Hence, this enzyme accepts the armless tRNAs to a certain extent and adds two C-residues, but not the terminal A, indicating that A incorporation requires a specific adaptation to such extreme substrates (Figure 2B, Figure 3, Table 1). In contrast, the *R. culicivorax* enzyme is adapted to these tRNA structures and efficiently adds the complete CCA-triplet. For *Hsa*CCA and *Rcu*CCA, the kinetic parameters for CC-incorporation on standard as well as armless tRNAs are quite similar and correspond to published values (Lizano *et al,* 2008). Yet, in both enzymes, these two reaction steps are adapted to structurally deviating tRNA substrates, whereas the *Eco*CCA is not able to catalyze this reaction. However, when it comes to A-addition, only *Rcu*CCA is adapted to the hairpin-like tRNAs (Figure 2B, Figure 3), although this reaction step is less efficient than C-addition (Table 1).

In the transition from C- towards A-addition, CCA-adding enzymes undergo a considerable domain rearrangement in order to accommodate the growing tRNA 3’-end in the catalytic core and to switch the specificity of the amino acid template (motif D) in the nucleotide binding pocket from CTP towards ATP recognition (Li *et al,* 2002; Toh *et al,* 2009; Ernst *et al,* 2015). During this structural rearrangement, the tRNA substrate has to remain bound to the enzyme, and this is usually accomplished by specific interactions of the tRNA 3’-end in the catalytic core and of the elbow region (T- and D-loop) with the C-terminal region of the CCA-adding enzyme (Tomita *et al,* 2004; Toh *et al,* 2009). For CCA-adding enzymes adapted to conventional tRNAs or tRNAs lacking only one arm, this interaction is not very tight, as no K_d_ values could be determined (Shi *et al,* 1998; Tretbar *et al,* 2011; Ernst *et al,* 2015) - yet it is sufficient for a complete synthesis of the CCA-end. For armless tRNA, however, these interactions seem to be insufficient, and the observed high substrate affinity of *Rcu*CCA (in contrast to the human enzyme; Figures 5, EV2) corroborates this hypothesis. Obviously, the *Romanomermis* enzyme is able to bind its substrate very tightly, and since this is also the case for armless tRNAs, this interaction cannot involve the conventional contacts between C-terminus of the enzyme and T-loop of the tRNA but must be located elsewhere, although the C-terminus contributes to the terminal A-addition to a certain extent, as shown by the *Rcu*/*Hsa*CCA chimera A (Figure 4).

In the detailed analysis of enzyme chimeras between *Rcu*CCA and *Hsa*CCA, a specific adaptation of the N-terminus is obvious. Here, two elements of the catalytic core are described to play an important role in the specificity switch from C- to A-addition. The flexible loop acts as a lever that adjusts the templating amino acids for correct ATP binding (Neuenfeldt *et al,* 2008; Just *et al,* 2008; Toh *et al,* 2009; Hoffmeier *et al,* 2010), and motif C represents a springy hinge that supports the domain rearrangements involved in this reaction step (Toh *et al,* 2009; Ernst *et al,* 2015). Accordingly, one could expect that both elements are adapted to the CCA-addition on armless tRNAs. Yet, chimera D (human enzyme with flexible loop of *Rcu*CCA) as well as chimera E (human enzyme with motif C of *Rcu*CCA) do not show any A-addition on the armless tRNA, while they are fully active on a standard substrate (Figure EV1). Structural data exclude a direct interplay between these elements (Toh *et al,* 2009; Ernst *et al,* 2015; Kuhn *et al,* 2015). Nevertheless, we analyzed chimera F, containing both elements of *Rcu*CCA, in case that the intervening region contributes to the function of these elements in A-addition. Yet, chimera F is also not able to incorporate the terminal A residue on the hairpin-like tRNA, so that a specific adaptation of the flexible loop and motif C can be excluded.

A further dissection of the N-terminal *Rcu*CCA components in the chimeras revealed that constructs carrying a β-turn element located between strands 3 and 4 of the β-sheet in the catalytic core are able to add the terminal A, indicating that this element represents a major adaptation to armless tRNAs (Figure 4, Figure EV1: chimeras B, C, G and H).

### The ß-turn element has an impact on the substrate affinity of *R. culicivorax* CCA-adding enzyme

In chimera H, the inserted β-turn element consists of 30 amino acids located between motif A and a conserved stretch upstream of the flexible loop (Figure 1). Toh *et al*. showed that in CCA-adding enzymes for conventional tRNAs, this region plays an important role in binding the growing CC-end of the tRNA and adjusting it in the catalytic core for the nucleophilic attack initiating A-addition (Toh *et al,* 2009). Similar β-turn elements are found for the archaeal class I CCA-adding enzymes (Tomita *et al,* 2006; Xiong & Steitz, 2004; Cho *et al,* 2005), poly(A) polymerases as well as for DNA- and RNA-polymerases, where they are described to position the 3’-hydroxyl of the primer 3’-end in close vicinity of the catalytically important metal ions located in the active site (Martin & Keller, 2004). In *Rcu*CCA, however, the β-turn element has an additional function, as it dramatically contributes to tRNA binding. While the turn itself (GEKH) is identical, the flanking sequences in the *Rcu*CCA enzyme differ in 13 positions from the counterpart. In these regions, the *Romanomermis* enzyme carries several lysine residues that are not present in *Hsa*CCA (Figure 6). While these residues probably contribute to the correct positioning of the tRNA 3’-end as a primer, they might also stabilize the interaction of the enzyme with its substrate.

**Figure 6.**
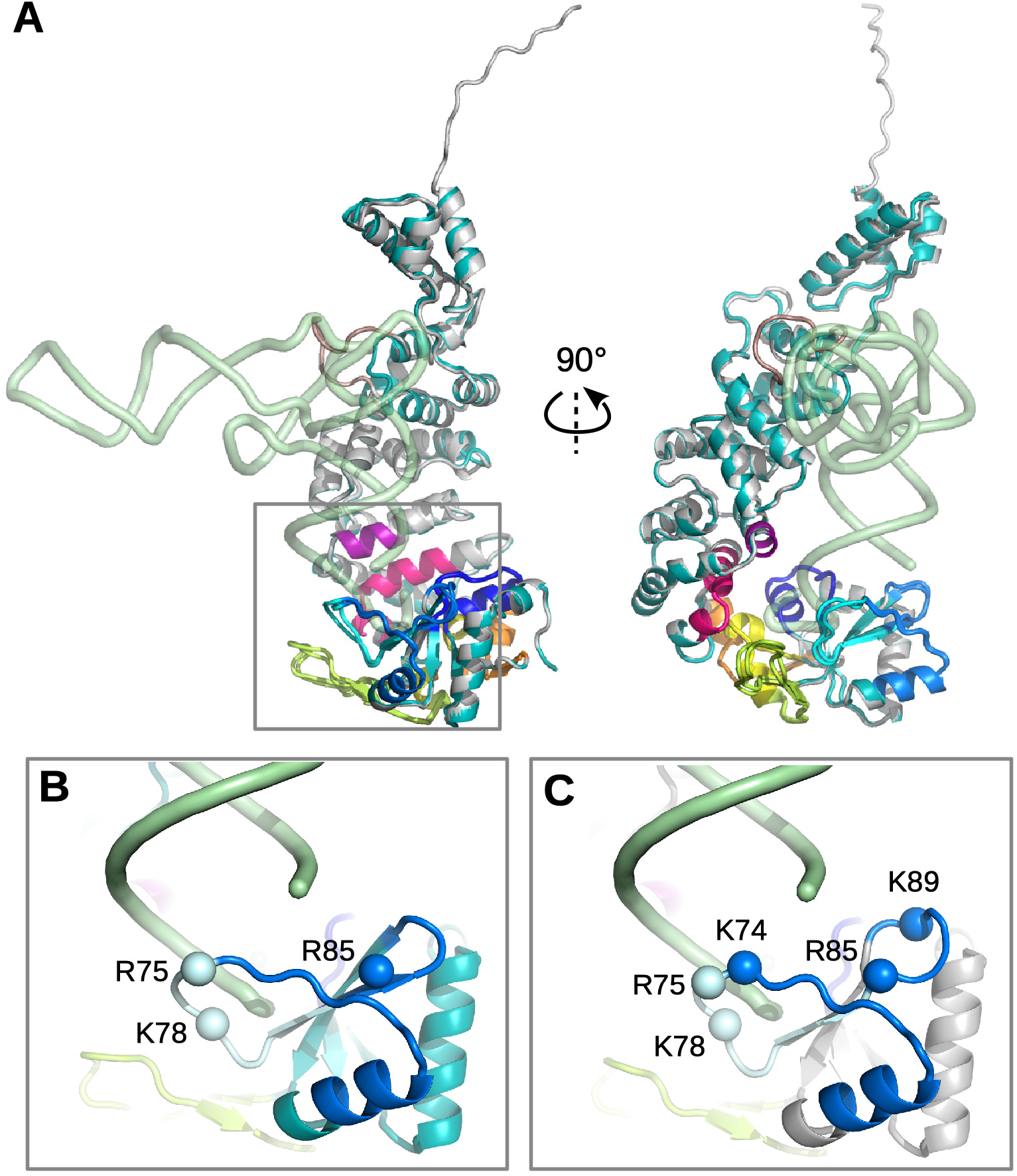
The β-turn in *Hsa*CCA and *Rcu*CCA enzymes. **A.** Superimposed full-length models of *Hsa*CCA (cyan) and *Rcu*CCA (light gray) with the transparent backbone of a bound tRNA (green; the tRNA position was obtained by superimposing the *T. maritima* complex onto the human enzyme) in two perpendicular views. Motif A (dark blue), β-turn region (medium blue), β-turn (light blue), B/A motif and flexible loop (green), motif B (yellow), motif C (orange), motif D (red) and motif E (violet) are indicated. **B, C.** Zoom into the β-turn region and the tRNA 3’-end (corresponding to the squared region in **A**) of *Hsa*CCA (**B**) and RcuCCA (**C**). Spheres represent the Cα positions of positively charged residues (K and R). *Rcu*CCA carries two additional lysines at positions 74 and 89 that might contribute to tRNA binding and primer positioning.

The presented results on *Rcu*CCA, *Hsa*CCA and the corresponding chimeras support the following hypothesis. As described above, the binding of the tRNA’s T-loop to its C-terminus represents one of the major substrate interactions of the CCA-adding enzyme, ensuring that the tRNA primer remains correctly located for A-addition during the domain rearrangements inducing the specificity switch. As this interaction is not possible with armless tRNAs, the human enzyme loses contact after adding two C residues, while a conventional tRNA remains correctly bound in the T-loop/C-terminus interaction and gets elongated by a complete CCA-terminus. In *R. culicivorax*, the CCA-adding enzyme had to adapt to armless tRNAs and evolved a different mode of tRNA interaction, as a binding of T-loop and C-terminus is not possible anymore. Here, the function of the β-turn region evolved from simple primer positioning for A-incorporation into an enhanced high-affinity substrate binding compensating for the loss of the conventional C-terminal tRNA interaction. This ensures that the enzyme’s interaction with armless tRNAs is sufficiently strong to survive the structural rearrangements during polymerization. A drawback of this tight binding might be a reduced product release, and the slightly less efficient CCA-addition on the conventional tRNA^Phe^ could be an indication of this (Table 1).

Taken together, the CCA-adding enzyme of *R. culicivorax* shows a remarkable adaptation to hairpin-like tRNA where the loss of substrate interactions with the C-terminus is compensated by enhanced tRNA binding of a different enzyme region. The evolutionary plasticity of enzymes is described for the composition of active site residues, where amino acids with identical catalytic roles are located at different positions in the primary sequence (Todd *et al,* 2002). The catalytic core, however, remains unchanged and structurally almost identical. In contrast, the β-turn element of *Rcu*CCA is not a mimicry of the T-loop/C-terminus interaction but recognizes a very different region of the tRNA (the 3’-end) and is still part of the catalytic core. Hence, its specific adaptation to the miniaturized tRNAs add a new layer of evolutionary plasticity. The structural resolution of this enzyme in complex with its tRNA substrate is expected to shed more light into this unusual and fascinating substrate adaptation.

## Materials and Methods

### Construction of recombinant enzymes

Open reading frames of CCA-adding enzymes from *Escherichia coli* and *Homo sapiens* were cloned into pET30 Ek/LIC plasmid with an N-terminal His_6_-Tag. The mt target signals were not included, and in both enzymes, the cloned coding regions started at the following conserved methionine residue as described (Reichert *et al,* 2001; Augustin *et al,* 2003; Lizano *et al,* 2007; Lizano *et al,* 2008). For the CCA-adding enzyme of *Romanomermis culicivorax*, the coding sequence was identified in the *R. culicivorax* genome assembly available at https://parasite.wormbase.org/Romanomermis_culicivorax_prjeb1358/Info/Index/, codon-optimized for expression in *E. coli* and synthesized in pET28a by GenScript^®^ (Piscataway, NJ, USA). All alignments were done using Jalview 2 (Waterhouse *et al,* 2009).

### Cloning of chimeric enzymes

Chimeric enzymes of CCA-adding enzymes from *Homo sapiens* and *Romanomermis culicivorax* were generated via site-directed mutagenesis in pET30-Ek/LIC or pET28a plasmids, respectively. All chimeras were cloned with an N-terminal His_6_-tag. The fusion positions of all chimeras are shown in Table EV1 and Figure EV3.

### Expression and purification of recombinant enzymes

*E. coli* BL21 (DE3) cca::cam lacking the endogenous CCA-adding enzyme were transformed with plasmids encoding the CCA-adding enzymes from *H. sapiens* (*Hsa*CCA), *R. culicivorax* (*Rcu*CCA) or chimeric enzymes. For CCA-adding enzyme from *E. coli* (*Eco*CCA), *E. coli* BL21 (DE3) was used. Cells were grown in 400 ml LB or TB with 50 μg/ml kanamycin and 35 μg/ml chloramphenicol (only for cca::cam strains) at 30°C. Expression was induced at OD_600_=1.5 by adding 400 ml ice-cold LB or TB containing both antibiotics and 2 mM IPTG to a final concentration of 1 mM. Cultures were incubated over night at 16°C and then harvested at 6,340 g for 15 min.

Pellets were resuspended in 8 ml ice-cold lysis-buffer (25 mM Tris/HCl pH 7.6, 500 mM NaCl, 1 mM DTT for *Eco*CCA and *Hsa*CCA, 100 mM phosphate buffer pH 7.0, 500 mM NaCl, 10 % glycerol, 0.2 % NP-40, 1 mM DTT for *Rcu*CCA) and disrupted with 5 g Zirconia beads and Fastprep-24 homogenizer (6 m/s, 30 s). Cell lysates were centrifuged at 30,600 g, 30 min, 4°C, sterile filtrated and loaded onto a HisTrap FF 1 ml or 5 ml column (GE Healthcare). Column wash was performed with 4-10 column volumes of binding buffer (25 mM Tris/HCl pH 7.6, 500 mM NaCl for *Eco*CCA and *Hsa*CCA, 100 mM phosphate buffer pH 7.0, 500 mM NaCl, 10 % glycerol for *Rcu*CCA) with 50 mM imidazole. His-tagged proteins were eluted with 3-8 column volumes of elution buffer (binding buffer with 500 mM imidazole). If necessary, protein containing fractions were further purified by size exclusion chromatography on a HiLoad 16/60 Superdex 75 pg column in binding-buffer containing 200 mM NaCl. Protein-containing fractions were combined and concentrated on Vivaspin 6 columns (15-30 kDa MWCO, GE Healthcare). Proteins were stored in 40 % glycerol (v/v) at − 80°C. Protein concentration was determined according to Bradford (Bradford, 1976).

### tRNA preparation

Armless mitochondrial tRNAs for isoleucine and arginine from *R. culicivorax* (Wende *et al,* 2014) and canonical cytosolic tRNA^Phe^ from *Saccharomyces cerevisiae* were generated as *in vitro* transcripts lacking the CCA-end in the presence of α^32^P-ATP (3000 Ci/mmol). In this preparation, homogeneous 5’- and 3’-ends of the transcripts were generated as described (Schürer *et al,* 2002). For kinetic analyses, tRNAs were transcribed without α^32^P-ATP.

### Electrophoretic Mobility Shift Assay (EMSA)

0.5 pmol α^32^P-ATP-labeled tRNA substrates were heated for 2 minutes at 90°C and incubated with 0 to 4 μM of enzyme in HEPES/KOH (pH 7.6), 30 mM KCl and 6 mM MgCl_2_ at 20°C for 10 minutes. After addition of 80% glycerol (final concentration of 18.5 %), tRNAs were separated by 5% native polyacrylamide gel electrophoresis. For visualization of enzyme-bound and free substrates, a Typhoon 9410 scanner was used (GE Healthcare). Dissociation constants were determined in three independent experiments by nonlinear regression using GraphPad Prism 7.

### Activity test and determination of arbitrary units

Enzyme activity was adjusted using canonical tRNA^Phe^ as a substrate. CCA-addition was performed in 30 mM HEPES/KOH pH 7.6, 30 mM KCl, 6 mM MgCl_2_, 0.5 mM NTP mix, 2 mM DTT. For calculation of arbitrary units, 5 pmol of tRNA were incubated with increasing amounts of enzymes for 30 min at 20°C. Reactions were ethanol-precipitated and analyzed on 10 % or 12.5 % polyacrylamide gels by autoradiography. Enzyme amounts leading to 50 % substrate-turnover were defined as 1 arbitrary unit. Nucleotide incorporation assays were performed as mentioned above. 5 pmol of tRNA were incubated with 1, 5, 10, 25 and 50 arbitrary units of enzyme for 30 min at 20°C and analyzed as described (Hoffmeier *et al,* 2010; Ernst *et al,* 2018).

### Kinetic analysis

Steady-state Michaelis-Menten kinetics were performed as described (Just *et al,* 2008). Each reaction contained 3 μCi of α^32^P-CTP or α^32^P -ATP (3000 Ci/mmol) and 30-35 ng *Hsa*CCA or 30-75 ng *Rcu*CCA. Non-labeled tRNA transcripts without CCA were titrated from 1-10 μM and incubated for 15-20 min at 20°C. Determination of incorporated radioactivity was performed as described (Just *et al,* 2008; Hoffmeier *et al,* 2010) and kinetic parameters of three independent experiments were calculated by non-linear regression Michaelis-Menten kinetic (GraphPad Prism). As the tRNA transcripts are not soluble at excessive saturating conditions, the calculated kinetic parameters represent apparent values, as frequently used for this type of enzymes (Tomita *et al,* 2004; Tomita *et al,* 2006; Hoffmeier *et al,* 2010; Wende *et al,* 2015; Ernst *et al,* 2018).

### Enzyme modeling

A homology model of *Rcu*CCA enzyme was built using Modeller (Webb & Sali, 2016) based on a sequence alignment with the human enzyme and on the human structure (PDB id: 4×4w) (Kuhn *et al,* 2015). A model of the human enzyme was also built to include all loops that are not visible in the crystal structure. Models were superimposed to the structure of *Thermotoga maritima* enzyme:tRNA complex (PDB id: 5hc9) (Yamashita & Tomita, 2016) in PyMOL (v2.4.0, Schrödinger) to position a tRNA in the active site and visualize loops in the vicinity of the tRNA 3’-tail.

## Acknowledgements

We thank Philipp Schiffer (University of Cologne) for help with the annotation of the CCA-adding enzyme from *Romanomermis culicivorax*. This work was supported by Deutsche Forschungsgemeinschaft Grant Mo 634/10-1.

## Author Contributions

HB, CS and MM designed research; SP, OH, TK, SB, KR and TJ performed research; HB, CS, MM, SP, KR and OH analyzed the data; OH, SP, HB and MM wrote the manuscript.

